# TCR Repertoire Analysis of CD4-Positive T Cells from Blood and an Affected Organ in an Autoimmune Mouse Model

**DOI:** 10.1101/2023.03.08.531799

**Authors:** Tatsuya Ishikawa, Kenta Horie, Yuki Takakura, Houko Ohki, Yuya Maruyama, Mio Hayama, Maki Miyauchi, Takahisa Miyao, Naho Hagiwara, Tetsuya J. Kobayashi, Nobuko Akiyama, Taishin Akiyama

**Affiliations:** Laboratory of Immune Homeostasis, RIKEN Center for Integrative Medical Sciences, Yokohama, Japan; Graduate School of Medical Life Science, Yokohama City University, Yokohama, Japan; Institute of Industrial Science, The University of Tokyo, Tokyo 153-8505, Japan

## Abstract

One hallmark of some autoimmune diseases is the variability of symptoms among individuals. Organs affected by the disease differ between patients, posing a challenge in diagnosing the affected organs. Although numerous studies have investigated the correlation between T cell antigen receptor (TCR) repertoires and the development of infectious and immune diseases, the correlation between TCR repertoires and variations in disease symptoms among individuals remains unclear. This study aimed to investigate the correlation of TCRα and β repertoires in blood T cells with the extent of autoimmune signs that varies among individuals. We sequenced TCRα and β of CD4^+^CD44^high^CD62L^low^ T cells in the blood and stomachs of mice deficient in autoimmune regulator (*Aire*) (AIRE KO), a mouse model of human autoimmune polyendocrinopathy-candidiasis-ectodermal dystrophy (APECED). Data analysis revealed that the degree of similarity in TCR sequences between the blood and stomach varied among individual AIRE KO mice and reflected the extent of T cell infiltration in the stomach. We identified a set of TCR sequences whose frequencies in blood might correlate with extent of the stomach manifestations. Our results propose a potential of using TCR repertoires not only for diagnosing disease development but also for diagnosing affected organs in autoimmune diseases.

## 1 Introduction

Autoimmune diseases result from disrupted immune systems that trigger inflammation against various organs. Some autoimmune diseases present highly heterogeneous symptoms, as the severity and affected organs vary among patients. For example, systemic lupus erythematosus can affect various organs, including joints, skin, kidneys, heart, the hematopoietic system, and the nervous system, but with varying degrees of involvement among patients (Allen, Rus, & Szeto, 2021; Wu et al., 2017). This individual variation is also seen in patients with APECED. APECED is caused by loss-of-function mutations in *AIRE*, a gene encoding the nuclear factor responsible for ectopic expression of peripheral-tissue antigens by thymic epithelial cells to mediate negative selection of self-reactive thymocytes in the thymus (Anderson et al., 2002; Perniola, 2018). Patients are diagnosed as APECED by developing at least two symptoms among endocrine deficiency, ectodermal dystrophy, and chronic mucocutaneous candidiasis (Betterle, Greggio, & Volpato, 1998; Perheentupa, 1996). While some organs, such as adrenal and parathyroid glands, are constantly affected, others like thyroid glands and the pancreas are affected in only 2-12% of cases, and gastritis is seen in 13-15% of cases (Betterle et al., 1998; Perheentupa, 1996).

Mice deficient in *Aire* (AIRE KO) were previously established and showed autoimmune phenotypes (Anderson et al., 2002; Goldfarb et al., 2021; Hubert et al., 2009; Jiang, Anderson, Bronson, Mathis, & Benoist, 2005; Niki et al., 2006; Ramsey et al., 2002; Su et al., 2008; Venanzi, Melamed, Mathis, & Benoist, 2008). Analysis of these mutant mice revealed that AIRE regulates thousands of tissue specific antigens (TSAs) in medullary thymic epithelial cells (mTECs). mTECs express and present these TSAs, thereby promoting negative selection of TSA-reactive T cells and generation of regulatory T cells in the thymus.

While autoimmune phenotypes in AIRE knockout (KO) mice were reported to be independent of microorganisms and their derivatives(Gray, Gavanescu, Benoist, & Mathis, 2007), the severity of symptoms, as well as the organs affected, can vary among different genetic backgrounds (Jiang et al., 2005; Niki et al., 2006), types of mutations (Goldfarb et al., 2021; Hubert et al., 2009; Kuroda et al., 2005; Su et al., 2008), and individuals (Anderson et al., 2002; Jiang et al., 2005; Ramsey et al., 2002; Venanzi et al., 2008). In addition to these deterministic factors, this variation may be ascribed partly to the stochastic nature of TSA expression regulated by AIRE(Dhalla et al., 2020; Meredith, Zemmour, Mathis, & Benoist, 2015), and to the inherent stochasticity that thymic T cells need to encounter mTECs, which are a rare population among total thymic cells, for thymic selection.

Because of the heterogeneous nature of autoimmune diseases, diagnosis of affected organs is necessary. In addition to the approach by detecting organ-specific protein markers (Suzuki et al., 2008; Zhang, Gao, Liu, & Zhang, 2023) (e.g. Neurofilament light chain in multiple sclerosis), it would be desirable to be able to use TCR repertoire analysis for diagnosis. TCR repertoire is formed by a combination of the hypervariable complementarity-determining region 3 (CDR3) loop of TCRs on their cell membranes and diverse V(D)J recombination in the thymus, allowing T cells to recognize myriad antigens in the body (Schatz & Ji, 2011). Since T cells play critical roles in development of autoimmune diseases by recognizing self-antigens, their TCR repertoires change depending on affected organs (Mitchell & Michels, 2020). Therefore, diagnosis by detecting their TCR repertoires prior to disease exacerbation may provide an earlier window for disease diagnosis than the approach detecting organ-specific protein markers For example, by using peripheral blood samples from patients, multiple studies have shown altered utilization of V and J genes in patients with systemic lupus erythematosus, rheumatoid arthritis, and Sjogren’s syndrome (Liu et al., 2019; Lu et al., 2022; Ye et al., 2020). Moreover, Moore et al. have performed TCR repertoire analyses of spleen, salivary gland, and brain choroid plexus from an MRL/lpr mouse model of lupus and identified a set of TCR sequences significantly detected in the brain (Moore et al., 2020). However, despite growing evidence suggesting a correlation between TCR repertories and development of autoimmune diseases, the correlation between TCR repertoires and variations in disease severity of autoimmune diseases has rarely been studied.

In the present study, we performed TCRα and β sequencing in the blood and stomachs of Aire-deficient mice (AIRE KO), a mouse model of APECED. The data suggest that some TCR repertoires in the blood and stomach were commonly increased in these mice and correlated with the extent of T cell infiltration in the stomach. This finding supports the diagnostic potential of TCR repertoire in monitoring the onset and affected organs of autoimmune diseases.

## 2 Results

### 2.1 Individual variation in phenotypes of AIRE KO mice

Our aim in this study was to seek the correlation of TCR sequence with the extent of autoimmune manifestations in a murine model. As a murine model of autoimmune diseases, we chose gastric autoimmunity of AIRE KO mice because: 1) the stomach is large enough to obtain sufficient numbers of T cells for TCR sequencing and 2) autoimmune inflammation in the stomach is heterogeneously induced in AIRE KO mice(Anderson et al., 2002; Jiang et al., 2005).

We first confirmed the heterogeneous features of autoimmune phenotypes in the stomachs of AIRE KO mice. Inflammatory status of the stomachs was evaluated for AIRE KO mice approximately 30 weeks old and their wild-type littermates (AIRE WT) as controls. Flow cytometric analysis indicated that the proportion and cell numbers of CD44^high^CD62L^low^ activated effector memory T cells (Teffs) among CD4^+^ and CD8^+^ T cells were significantly increased compared to controls (Fig. 1a-b and Fig. S1), manifesting stomach inflammation with activated T cell infiltration in mutant mice. In contrast, the frequency of Foxp3^+^CD25^+^ was not changed in the stomach and blood of AIRE KO mice compared to AIRE WT (Fig. S2).

**Figure 1.**
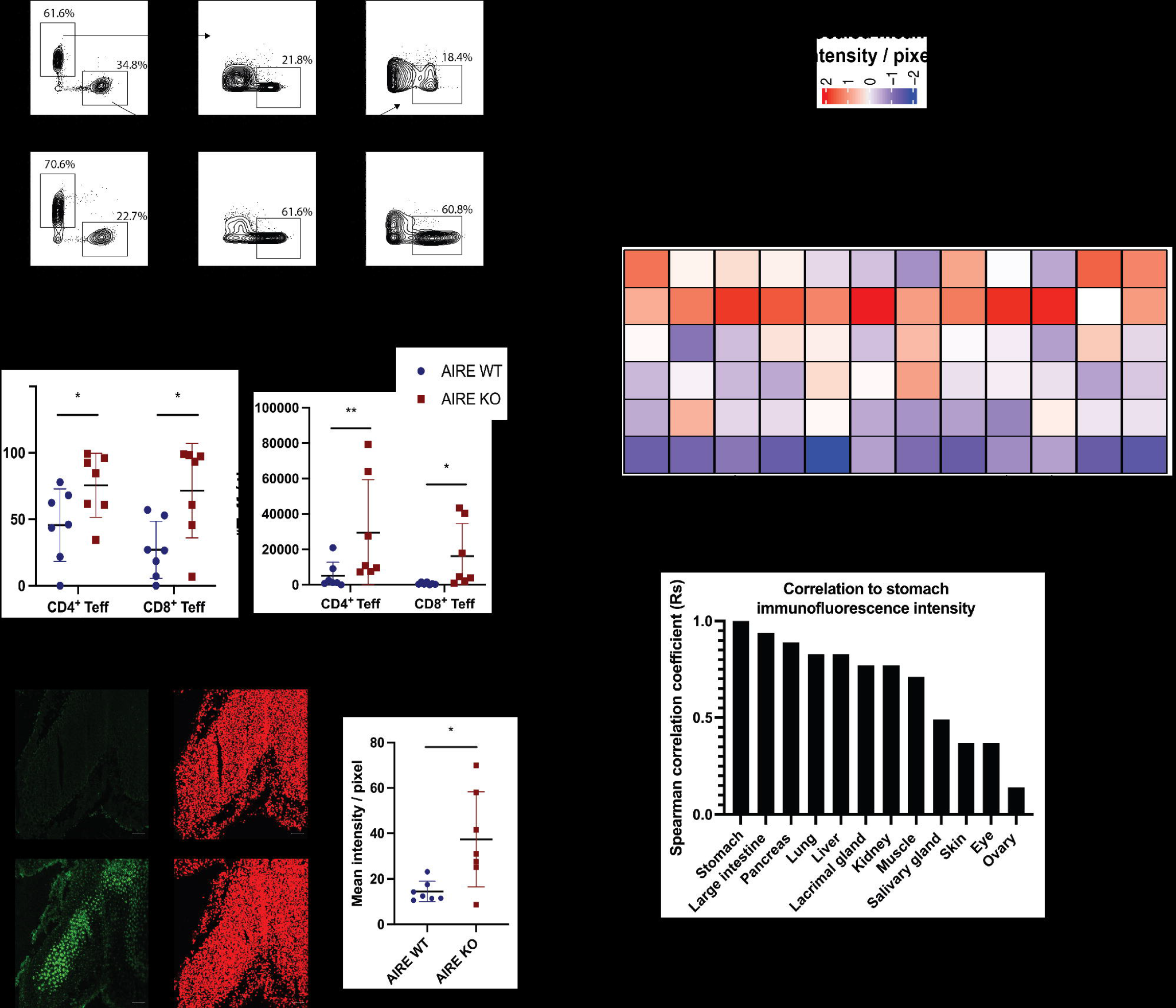
Individual variation in multi-organ phenotypes of AIRE KO mice. (a) Flow cytometric analysis of CD4^+^ and CD8^+^ Teffs from stomachs of AIRE WT and AIRE KO mice. (b) Numbers and percentages of CD4^+^ and CD8^+^ Teffs from stomachs of AIRE WT and AIRE KO mice. Two-tailed Student’s t-test and Mann-Whitney U test were used for statistical analyses of the cell number and the percentage, respectively. (c) Immunofluorescent staining of AIRE WT and AIRE KO sera against stomach sections of *Rag1*^-/-^ mice. Data were analyzed using Mann-Whitney U test. (d) Heatmap of scaled mean intensity of immunofluorescent signals. (e) Spearman correlation coefficients (Rs) between stomach immunofluorescence signals and those of other organs. ns, not significant (p >0.05). *p < 0.05, **p < 0.01.

We further evaluated generation of autoantibodies against stomach tissues by immunohistochemical staining of the stomach section from RAG1-deficient (*Rag1*^−/−^) mice with serum of AIRE KO mice (Fig. 1c, left). Quantification of these immunofluorescent images showed that the level of autoantibodies against the stomach was significantly higher in the serum of AIRE KO mice than that of AIRE WT mice (Fig. 1c, right), which is consistent with previous studies (Gavanescu, Kessler, Ploegh, Benoist, & Mathis, 2007). Notably, standard deviations (SD) in the number of CD4^+^ Teffs and serum levels of anti-stomach antibodies are higher in AIRE KO mice than in controls (Fig. 1a-c, Standard deviation (SD) = 27780 and 7076 each for the CD4^+^ Teff number and SD = 17061 and 614 for the CD8^+^ Teff number in AIRE KO and control mice, respectively; SD = 19.3 and 4.17 each for the serum antibody level in AIRE KO and control mice, respectively). Thus, there was large variation among individual AIRE KO mice in the number of infiltrated Teffs in the stomach and serum levels of anti-stomach antibodies. Consequently, significant individual variation in stomach manifestations was confirmed in this line of AIRE KO mice.

Dysfunction of AIRE provokes autoimmunity targeting multiple organs. We next investigated how the extent and target organs of autoimmunity vary in this Aire-deficient mice line from individual to individual. Titers of autoantibodies to various organs in the serum were quantified by immunohistochemical staining (Fig. S3). These results showed that the profile of targeted organs varied markedly among individual mice (Fig. 1d-e). Mouse #1 was the most severely affected, as it produced the broadest profile with the greatest number of organs having high autoantibody signal intensity. On the other hand, Mouse #6 showed a sporadic profile with the highest signal intensity for the stomach, lung, and pancreas. Mice #3, #4, and #5 also produced sporadic profiles with relatively high intensity for organs such as skin, eyes, and salivary glands. Finally, Mouse #2 produced the smallest signal in the greatest number of organs examined. We also evaluated the correlation of autoantibody signal intensity of stomach with that of other organs. While organs such as large intestine, pancreas, and lung showed high correlation, skin, eyes, and ovaries showed low correlation with stomachs, suggesting that titers of serum autoantibodies against each tissue vary among individuals in these AIRE KO mice (Fig. 1d and e). Taken together, these results indicate that autoimmune phenotypes of AIRE KO mice used in this study are heterogeneous. Accordingly, AIRE KO mice can comprise a model to assess the potential of TCR repertoires to monitor disease onset and affected organs.

### 2.2 Limited correlation of blood T cell count, TCR diversity, clonality, CDR3 length, and V/J usage with disease severity in AIRE KO mice

Before performing TCR sequencing, we asked whether blood T cell counts might reflect the onset of autoimmunity in AIRE KO mice. To this end, we used flow cytometry to detect CD4^+^ and CD8^+^ Teffs in the blood of AIRE KO and AIRE WT mice. We found that both the proportion and the number of CD4^+^ and CD8^+^ Teffs were comparable in AIRE KO and AIRE WT mice (Fig. 2a-b). Thus, the Teff count in blood does not reflect the onset of autoimmunity in AIRE KO mice.

**Figure 2.**
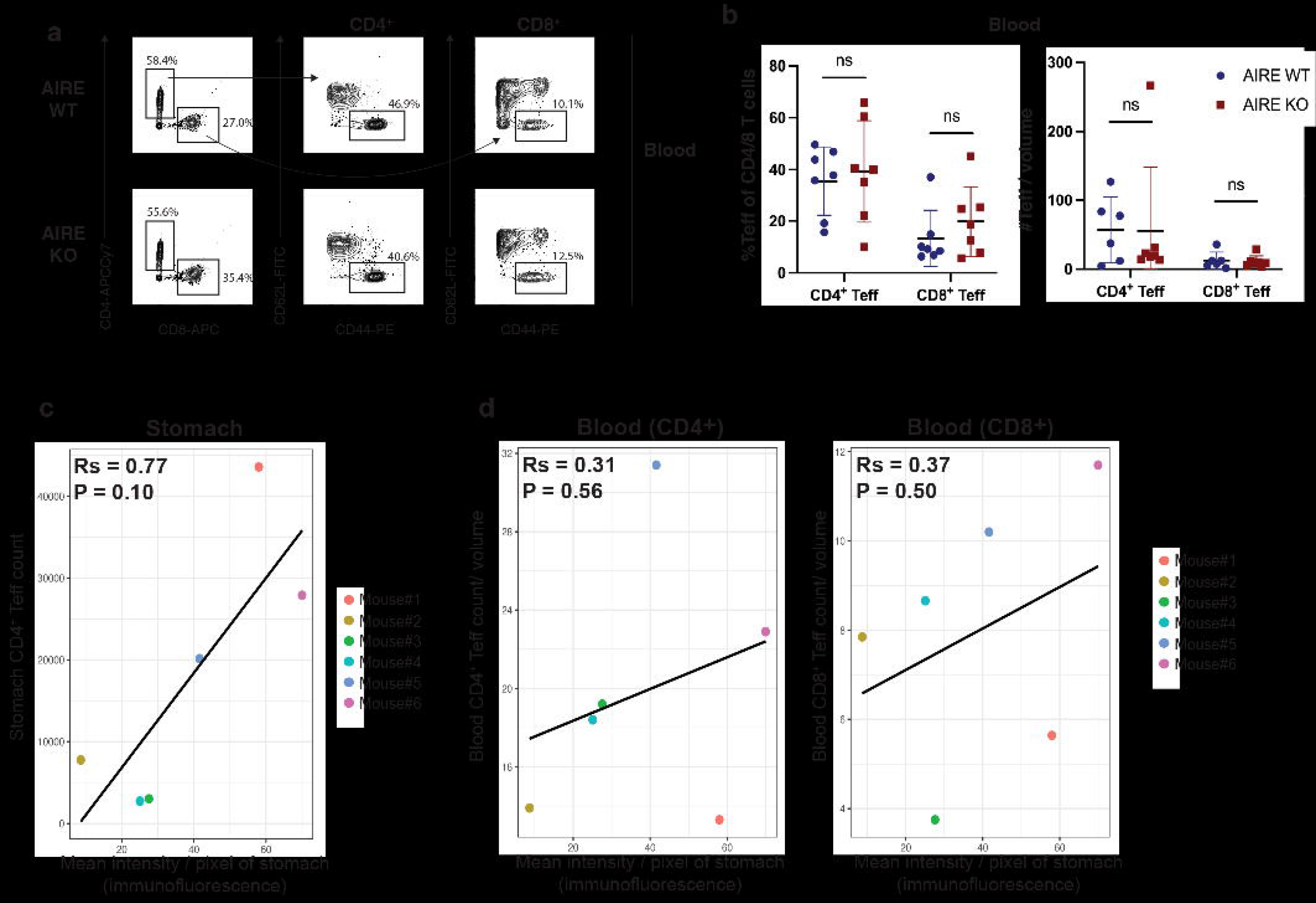
Correlation of activated T cell number in the stomach and blood with titer of anti-stomach antibody in AIRE KO mice. (a) Flow cytometric analysis of CD4^+^ and CD8^+^ Teffs from the blood of AIRE WT and AIRE KO mice. (b) Numbers and percentages of CD4^+^ and CD8^+^ Teffs from the blood of AIRE WT and AIRE KO mice. Two-tailed Student’s t-test and Mann-Whitney U test were used for statistical analyses of the cell number and the percentage, respectively. (c) Spearman’s correlation of stomach CD4^+^ Teff count with mean signal intensity of stomach immunofluorescence, reflecting the titer of anti-stomach antibody. (d) Spearman’s correlation of blood CD4^+^ Teffs (left) or CD8^+^ Teffs (right) with the mean signal intensity of stomach immunofluorescence. Rs, Spearman’s correlation coefficient. P, p value determined from Spearman’s correlation.

For TCR sequencing analysis, we sorted CD4^+^ Teffs in the blood and stomach (Table S1) because autoimmune phenotypes of AIRE KO are predominantly B-cell mediated (Gavanescu, Benoist, & Mathis, 2008). Approximately 5000 to 15000 CD4^+^ Teffs were FACS-sorted from the blood of WT (Blood WT) and AIRE KO (Blood KO). For the stomach, all CD4^+^ Teffs present in the organ were sorted. The total number of CD4^+^ Teffs obtained from the AIRE KO stomach (Stomach KO) ranged from 2000 to 40000 per stomach (hereafter referred to as stomach Teff count). The CD4^+^ Teff number in the WT stomach was too low to be used for TCR sequencing. Total RNA of these sorted CD4^+^ Teffs was prepared successfully from 6 AIRE KO (labeled from #1-6) and 7 AIRE WT (labeled from #7-13) mice with sufficient quality for TCR sequencing. Of note, the stomach Teff count showed a weak correlation with the fluorescence intensity of stomach immunostaining, which reflects the titer of autoantibodies in the serum (Fig. 2c, Rs = 0.77, P = 0.10). In contrast, as expected, there was no correlation between the number of blood CD4^+^ (Rs = 0.37, P = 0.56) or CD8^+^ Teffs (Rs = 0.37, P = 0.50) with the mean signal intensity of stomach immunofluorescence (Fig. 2d).

We performed TCR repertoire sequencing of these sorted CD4^+^ Teffs (Table S1). To consider the difference in cell number, the amount of RNA extracted from each sample was normalized for cDNA library preparation. By analyzing sequencing data, some parameters of TCRα and β repertoires such as diversity, clonality, CDR3 length, and V/J usage, were determined in each sample. We then sought to find parameters that predict disease onset in the stomachs of these AIRE KO mice according to two criteria: 1) the difference between Blood KO and Blood WT, for prediction by measurement of blood TCR repertoires, and 2) the similarity between Blood KO and Stomach KO, for supporting the reflection of stomach manifestations.

First, we calculated the number of unique clonotypes and normalized Shannon indices to evaluate diversity and clonality, respectively, of TCRs from Blood WT, Blood KO, and Stomach KO. We found that diversity among Blood WT, Blood KO, and Stomach KO was comparable for both TCRα and β (Fig. 3a). Chao1, another diversity index, was also comparable among all groups for both TCRα and TCRβ (Fig. S4). On the other hand, we found that TCRα clonality was significantly increased in Stomach KO in the Shannon index (Fig. 3a) while TCRβ clonality was comparable among all groups. Consistently, the Inverse Simpson index was high for TCRα in Stomach KO (Fig. S4) and that of TCRβ was relatively comparable among all groups. Overall, the diversity is comparable in both TCRα and TCRβ in AIRE KO whereas TCRα clonality of Stomach KO is higher as compared to that of blood samples.

**Figure 3.**
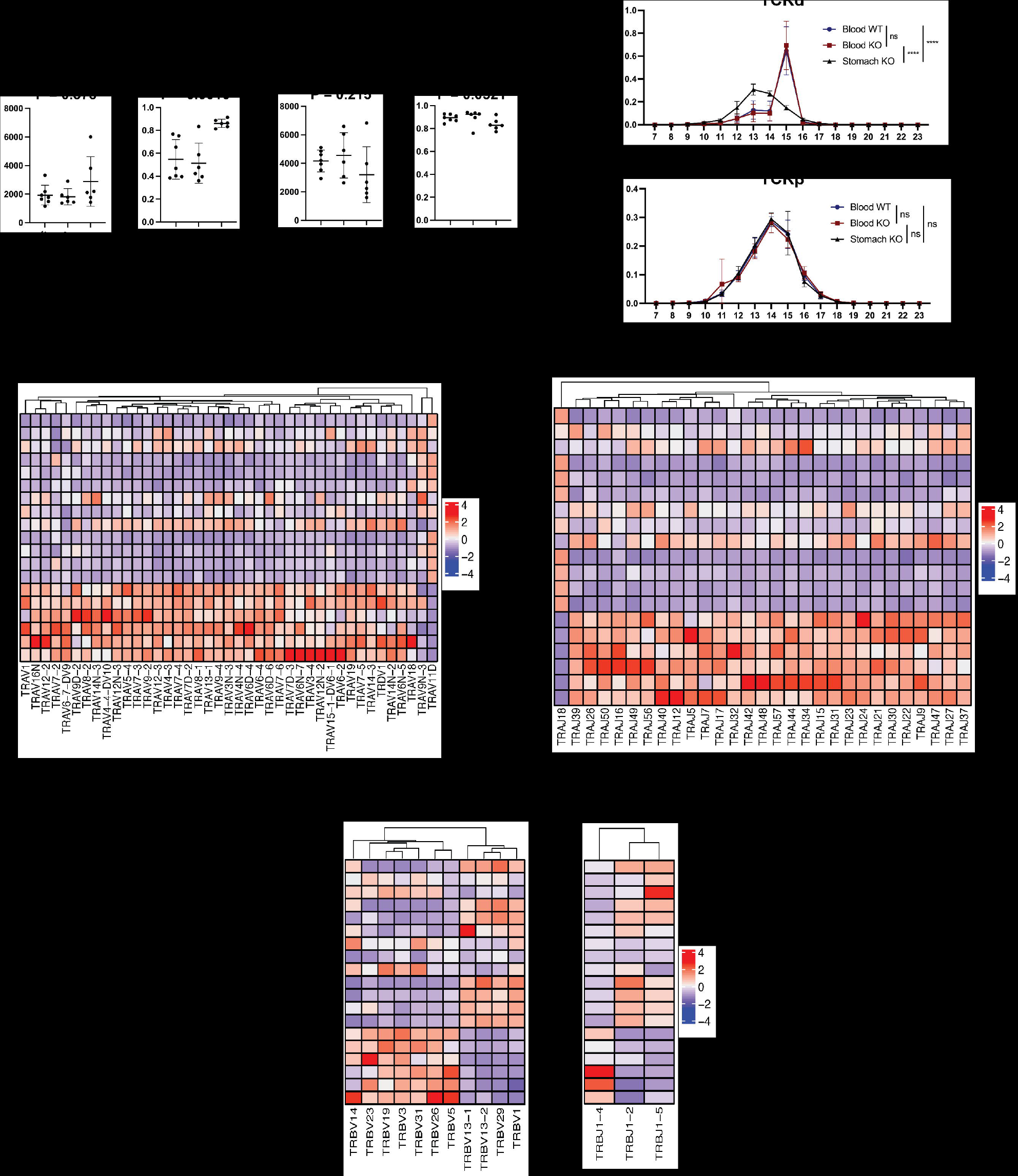
TCR repertoire analysis of blood and stomachs of AIRE KO mice. (a) Diversity and clonality of TCRα and β of Blood WT, Blood KO, and Stomach KO. Kruskal-Wallis test was used for comparisons and its p value, P, is shown. (b) CDR3 amino acid length distributions of TCRα and β of Blood WT, Blood KO, and Stomach KO. Two-way ANOVA with random variables. ****p < 0.0001. ns, not significant (p > 0.05). Heatmap showing scaled frequencies of significantly detected Vα (c), Jα (d), Vβ, and Jβ (e) genes between Blood WT and Blood KO and between Stomach KO and Blood WT/KO.

Similar to clonality, the amino acid length of CDR3 significantly decreased in Stomach KO for TCRα but remained comparable among all groups for TCRβ (Fig. 3b). Similar CDR3 length distributions following a Gaussian pattern were also observed in T cells from diseased organs in other mouse models of autoimmune disease (Moore et al., 2020). Although there were slight changes in amino acid usage at each position of CDR3 in Stomach KO compared to blood samples, no clear trend emerged (Fig. S4c). In support of the trend of clonality and CDR3 length, most significantly detected V and J genes were strongly present in Stomach KO for TCRα, but were comparably present in all groups for TCRβ (Fig. 3c-e). Since diversity, clonality, CDR3 length, and V/J usage for TCRα and β repertoires did not meet criteria for correlation, we concluded that these measures do not reflect disease severity in AIRE KO.

### 2.3 Individual similarities of TCR repertoires in the blood and stomachs of AIRE KO

We then shifted our attention to individual sequences in TCRα and β chains and focused on shared TCR sequences of CD4^+^ Teffs in the blood and stomach from the same AIRE KO mice. To this end, we calculated Jaccard indices to evaluate the similarity of TCRα and β repertoires based on the proportion of shared TCR sequences between each paired blood (Blood#1-13) and stomach sample (Stomach #1-6). Scatterplots and heat maps of Jaccard indices suggested that Blood KO correlated with the corresponding Stomach KO (Fig. 4a, b, and Table S2 and S3). Indeed, Jaccard indices of the blood and stomach from the same AIRE KO mice (on average about 4%) were four times higher than those of blood and stomachs from different mice (on average about 1%) for both TCRα and β (Fig. 4c). Thus, TCRα and β repertoires in the blood correlate with those in the stomach repertoire of AIRE KO mice, and more importantly, the correlation of each TCR repertoire between stomach and blood is individualized. Notably, scaled Jaccard indices correlated with the stomach Teff count, a measure reflecting the severity of the stomach manifestation (Fig. 4d, P =0.058 for TCRα and P = 0.0024 for TCRβ). Thus, mice with more severe gastric inflammation may have more common TCR repertoires in blood and stomach. Overall, these data support the potential for monitoring the severity of stomach manifestation by detecting TCR sequences common to the blood and stomach in AIRE KO.

**Figure 4.**
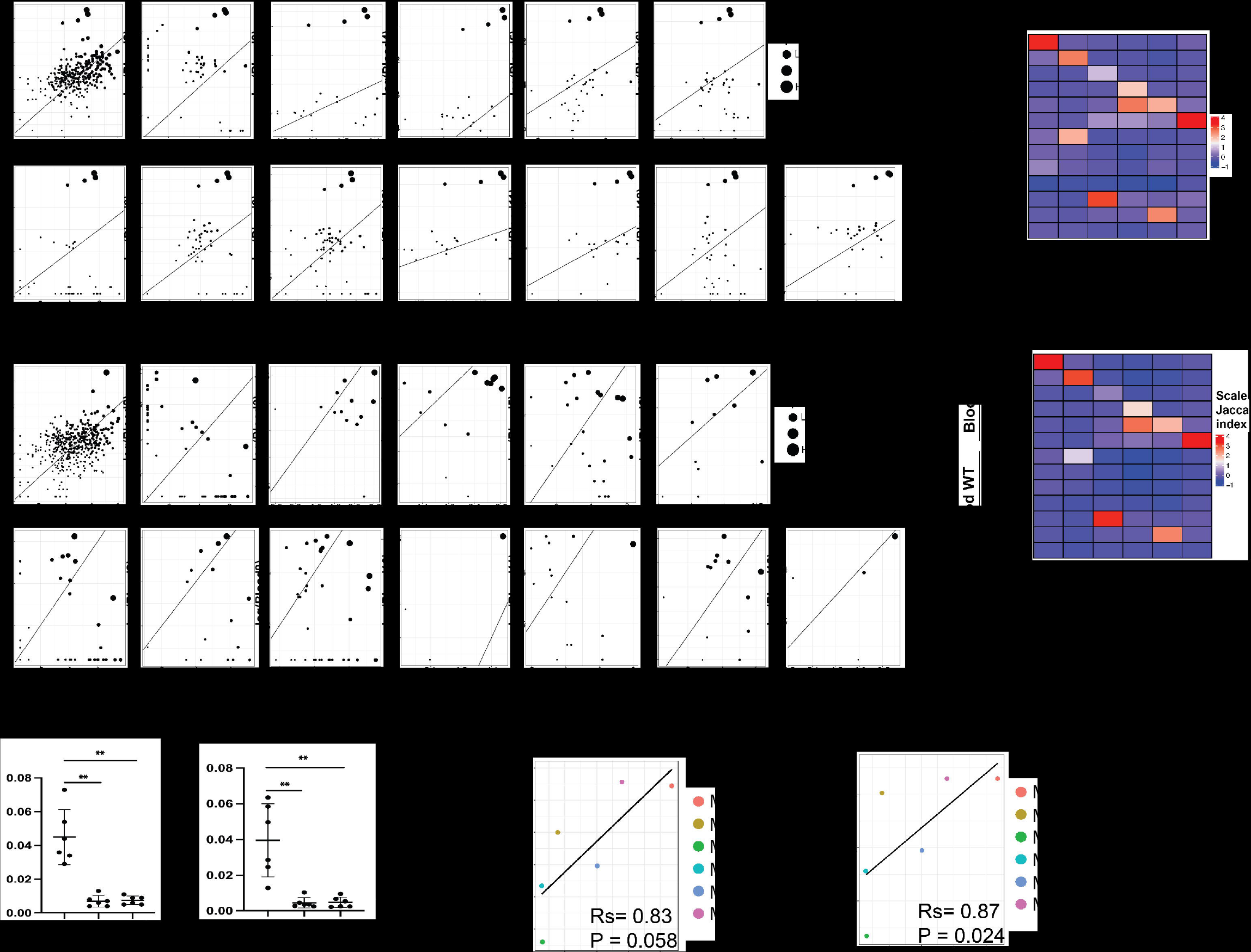
Individual similarity of TCR repertoires between blood and stomachs of AIRE KO. (a) Scatterplot showing frequencies of TCRs shared between Stomach 1 and each of the blood samples. (b) Heatmap of scaled Jaccard indices of each of blood and stomach pair. (c) Self-Blood KO vs Stomach: Jaccard indices of the stomach and blood from the same AIRE KO mice are plotted. Nonself-Blood KO vs Stomach: For each AIRE KO mice, average Jaccard indices of the stomach and blood from different AIRE KO mice are plotted. Nonself-Blood KO vs Stomach: For each AIRE KO mice, average Jaccard indices of the stomach and blood from AIRE WT mice are plotted. Mann-Whitney U test. **p < 0.01. (d) Spearman’s correlation between stomach Teff counts and scaled Jaccard indices of the blood and stomach from each AIRE KO mouse (from #1 to #6). Rs, Spearman’s correlation coefficient.

### 2.4 Identification of candidate TCRα and β repertoires to monitor disease severity in AIRE KO

We next sought to determine sequences of TCRα and β repertoires in CD4^+^ Teffs that correlate with Stomach CD4^+^ Teff count. Such TCR repertoires could monitor the severity of stomach manifestations in AIRE KO mice as a biomarker. To this end, we first screened the CDR3 of TCR sequences according to three criteria: 1) CDR3 sequences that were detected in both the Stomach KO and Blood KO of the same mice, 2) were found in at least two AIRE KO mice and 3) were not present in Blood WT. As a result, we successfully identified 29 CDR3 sequences for both TCRα and β. We then categorized these CDR3 sequences based on their amino acid sequences using the tcrdist3 package to assign mismatch distance between two CDR3 sequences based on the BLOSUM62 substitution matrix (Dash et al., 2017). The matrix gives the score that determines whether an amino acid substitution is conservative or nonconservative, reflecting the similarity of biochemical properties of two amino acids. The distance matrix was then used for hierarchical clustering. Based on the clustering result, we categorized identified CDR3 amino acid sequences into 4 clusters (k-means = 4) for TCRα (Fig. 5a) and 3 clusters (k-means = 3) for TCRβ (Fig. 5b).

**Figure 5.**
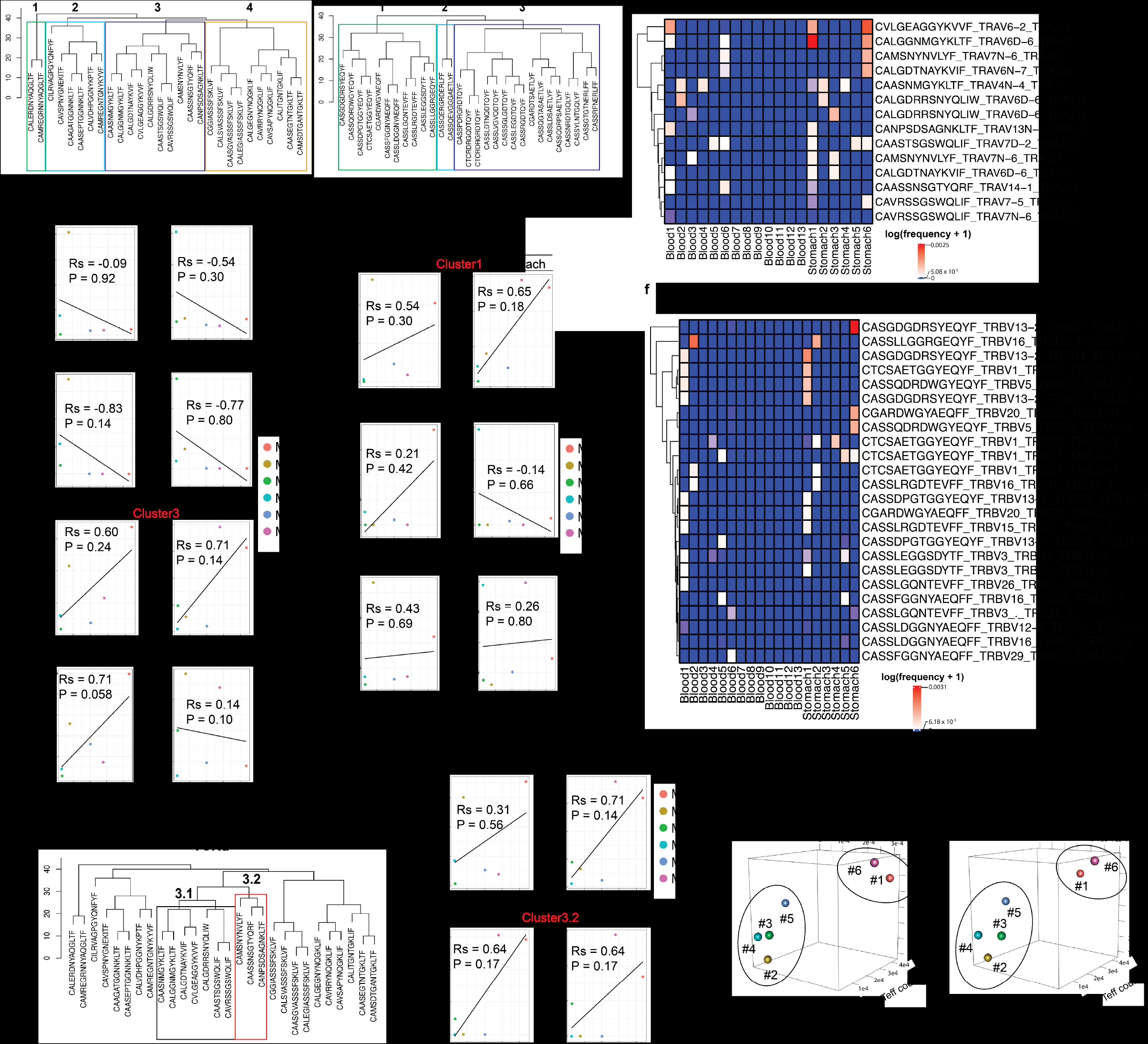
Identification of candidate TCR repertoires correlating with severity of stomach manifestation in AIRE KO. Hierarchical clustering of candidate TCRα (a) and β (b) repertories. Cluster numbers are shown at the tops of trees. Spearman’s correlation of stomach CD4^+^Teff counts and mean frequencies of TCRα (c) and β (d) sequences contained in each cluster detected in the blood (left) and stomach (right) of each AIRE KO mouse. Heatmap showing log-transformed frequencies of TCRα sequences contained in Cluster3 (e) and TCRβ sequences contained in Cluster1 (f). Different CDR3 nucleotide sequences encoding the same CDR3 amino acid sequences are also shown in the heatmap. (g) Hierarchical clustering tree showing subclusters of Cluster3 of TCRα (Cluster3.1 and 3.2). (h) Spearman’s correlation of stomach CD4^+^Teff counts and mean frequencies of TCRα sequences in Clusters 3.1 (top) and 3.2 (bottom) detected in the blood (left) and stomach (right) of each AIRE KO mouse. (i) 3-dimensional plots of mean frequencies of TCRα sequences in Cluster 3.2 detected in the blood (left) or stomach (right), stomach CD4^+^Teff counts, and mean signal intensities of stomach immunofluorescence. Rs, Spearman’s correlation coefficient.

Since a biomarker should discriminate between diseased and non-diseased subjects, we searched for clusters containing CDR3 sequences whose frequencies from the blood and stomach correlated with the severity of stomach manifestations, represented by the stomach Teff count. We found that the mean frequencies of CDR3 sequences classified as Cluster3 in TCRα and classified as Cluster1 in TCRβ (hereafter referred to as Cluster3/TCRα and Cluster1/TCRβ, respectively) showed some positive correlation with both the blood and stomach (Rs > 0.5) (Fig. 5c-d and Fig. S5). Stomach TCR frequencies of Cluster 3/TCRα and Cluster1/TCRβ were able to discriminate the two mice with the highest stomach Teff counts (Mouse #1 and Mouse #6) from the other mice. However, blood TCR frequencies of these clusters were not clearly correlated with stomach Teff counts (Fig. 5c-f). Especially for Cluster1/TCRβ, high TCR frequencies in the stomach of Mouse #6 were not reflected in the blood (Fig. 5d, f). Therefore, we focused on Cluster3/TCRα and found that when we divided Cluster3 into two sub-clusters, Clusters 3.1 and 3.2 (Fig. 5g), TCR frequencies of Cluster 3.1 were relatively high in the blood of Mouse #2 (Fig. 5e, h, and Table S4), a mouse with a low stomach Teff count (Fig. 5h). By contrast, TCR frequencies of Cluster 3.2 showed a relatively strong correlation with stomach Teff counts for both the blood (Rs = 0.64) and stomach (Rs = 0.64) (Fig. 5h). Therefore, we used the TCR frequencies of Cluster 3.2 from the blood or stomach and finally confirmed their correlations with two distinct dimensions reflecting the severity of stomach manifestations, stomach Teff count and mean signal intensity in immunofluorescent staining of the stomach (Fig. 5i). The result of 3D plots showed that by using TCR frequencies of Cluster 3.2 from both the blood and stomach, Mouse #1 and Mouse #6 are clearly distinguished from the others.

## 3 Discussion

In the present study, we used an AIRE KO mouse model of APECED to test whether TCR repertoires can serve as biomarkers to monitor organ manifestations in autoimmune disease. We proposed a set of TCR sequences in Cluster 3.2 of TCRα that may discriminate mice with relatively severe autoimmunity from other mice. These data support the potential of TCR repertoires to monitor disease severity, at least in this AIRE KO mouse model of autoimmune disease. However, it should be noted that there were relatively large individual differences in the experimental data for CD4Teff in the stomach and blood. These differences may be due to inherent variation among individuals, as well as slight differences in the rearing environment, such as food accessibility. To address these issues, it will be crucial to collect data from a larger number of samples and to analyze T cells obtained from additional organs in the future.

A previous study suggested that hydrophobic amino acid residues are enriched in CDR3 region of the TCRβ chain in self-reactive T cells (Stadinski et al., 2016). However, we did not observe a similar trend among all samples, including Stomach KO, which would be expected to have an enrichment of activated self-reactive T cells. Instead, our data revealed significant alterations in the clonality, CDR3 length, and V/J usage of Stomach KO repertoires for TCRα as compared to those of Blood WT and Blood KO, while the changes in TCRβ repertoires were less pronounced. The reason for the preferential change in TCRα over TCRβ should be clarified in the future. On the other hand, we found no significant differences in diversity, clonality, CDR3 length, or V/J usage between Blood WT and Blood KO. This aligns with a previous study demonstrating comparable TCR repertoire diversity in the blood of AIRE WT and AIRE KO mice (Oftedal et al., 2017).

To date, most analyses of TCR repertoire in autoimmune diseases have been conducted to monitor disease development rather than variation in disease manifestations (Mitchell & Michels, 2020). Most previous studies simply compared TCR repertoires between patients and normal individuals and searched for common TCR sequences that were abundant in patients (Liu et al., 2019; Lu et al., 2022; Ye et al., 2020). In addition to our current study, further investigations would be necessary to address the potential of TCR repertoires to dissect disease heterogeneity.

Despite the identification of candidate TCR repertoires to monitor stomach manifestation in AIRE KO mice, there is no guarantee that these TCR repertoires specifically recognize a self-antigen derived from the stomach. To address this issue, it is important to determine TCR repertoire specificities against stomach-derived antigens. However, a significant limitation of our approach lies in the fact that we conducted bulk-RNA-seq analysis separately for TCR α and TCR β components. As a result, we are unable to ascertain which specific TCR α and β pair is responsible for recognizing antigens in the stomach. Moreover, we could not find potential target epitopes of the identified TCR sequences in Cluster 3.2 by searching VDJdb, a curated database of TCRs with known antigen specificities (Shugay et al., 2018). Previously, MUCIN6 has been reported as a self-antigen involved in development of autoimmunity in AIRE KO mice(Gavanescu et al., 2007), and it is of interest to know whether identified TCR repertories can respond to MUCIN6. However, determining the antigen specificity of TCRs based on current experimental approaches is time-consuming, and there is a significant lack of data on known TCR-peptide pairs to develop a reliable algorithm for predicting antigen specificity of any TCR (Hudson, Fernandes, Basham, Ogg, & Koohy, 2023). Thus, it is imperative for future research to employ high-throughput experimental approaches that can concurrently identify numerous TCR-peptide pairs, exemplified by the high-resolution analysis offered by single-cell TCR repertoire analysis. Such experiments are pivotal in ensuring accurate predictions of tissue specificity within TCR repertoires.

The advantage of using a mouse model of autoimmune disease is that unlike human samples, it is easy to obtain samples from blood and peripheral organs. Especially for the purpose of this study, it would be very difficult to obtain sufficient T cells from stomachs of patients with autoimmune diseases for TCR repertoire analysis. However, TCR sequences of mice and humans are different. Clearly, there is a great need for bioinformatics tools that can predict human TCR sequences from those of mice, so that we can seamlessly translate findings from animal models to the clinic.

## 4 Experimental procedures

### 4.1 Mice

Female littermates were used in these experiments. Aire KO mice on the background of C57BL/6 were originally from the RIKEN RBC through the National Bio-Resource Project of MEXT in Japan (RBC03515: B6.Cg-Aire<tm2Mmat>/Rbrc) mice. B6.129S7-Rag1tm1Mom/J (*Rag1*^*–/–*^) were purchased from Jackson Laboratory. All mice had free access to food prior to the study. All mice were maintained under pathogen-free conditions and handled in accordance with Guidelines of the Institutional Animal Care and Use Committee of RIKEN, Yokohama Branch (2018-075).

### 4.2 Cell preparation and flow cytometry

When at least one pair of WT and KO were born from the same female mouse, the mouse pair were served for analysis when they reached 30 weeks of age. Stomach tissue was excised and immediately washed with RPMI medium. It was then minced before digestion with 1.0mg/mL collagenase in RPMI medium (Wako) at 37°C. Supernatant containing cells was then passed through a 40-μm filter. Cells were isolated from blood by adding 2 mL of RBC lysis buffer (Biolegend) per 100 uL of blood collected. After incubation with RBC lysis buffer for 10-15 min at room temperature, cells were then washed with FACS buffer before antibody staining. For flow cytometry, cells were first incubated with blocking antibodies against Fc receptors (Biolegend) before staining with antibodies against TCRβ, CD4, CD8, CD44, and CD62L. Dead cells were excluded by staining with 7-aminoactinomycin D. For detection of Tregs, intracellular staining was performed by incubating cells with fixation buffer for 30 min and washing with permeabilization buffer (Invitrogen) prior to incubation with blocking antibodies and antibodies against Foxp3. Cells were sorted using a FACS Aria instrument (BD) for TCR sequencing.

### 4.3 TCR library preparation and sequencing

Total RNA was prepared using RNeasy Micro Kit (QIAGEN) according to the manufacturer’s protocol. RNA samples with a RIN value of 8 or higher quality in the Bioanalyzer system were used for TCR repertoire analysis (7 mice for WT and 6 mice for KO). A cDNA library for TCR was prepared from RNA using a SMARTer Mouse TCR a/b Profiling Kit (Takara Bio) according to the manufacturer’s protocol. To account for differences in numbers of sorted cells, RNA quantities were normalized prior to cDNA preparation. cDNA libraries of both TCRα and β were then subjected to paired-end sequencing using a Miseq (Illumina) according to the protocol for Miseq Reagent Kit v3 (Illumina).

### 4.4 TCR data processing and analysis

TCRα and β sequencing data were mapped using MiTCR software (Bolotin et al., 2013). Output files from MiTCR were then converted to a format compatible with VDJtools, an analysis framework for repertoire sequencing data (Shugay et al., 2015). Diversity, clonality, CDR3 length, and significant V/J usage were calculated using VDJtools command lines. The Jaccard index of TCR repertoires between two samples was calculated by dividing the number of shared unique TCR sequences by the total number of unique TCR sequences. The distance matrix of selected TCR sequences was generated using the tcrdist3 python package (Dash et al., 2017), and hierarchical clustering was performed using the hclust function from the stats package in R.

### 4.5 Immunofluorescence

*Rag1*^-/-^ tissues were snap-frozen in OCT compound. Tissues were sectioned with a cryostat (6 µm) and fixed in cold acetone for 10 min. After washing with PBS, sections were blocked with 10% normal goat serum. Sera from AIRE KO mice (100x) and control (100x) were then applied to the tissue sections and incubated for 1 h. Secondary anti-mouse antibody conjugated to Alexa-488 dye was then added along with 1 µg/mL propidium iodide (PI) and 5 µg/mL RNase A and incubated for 40 min. All images were captured using an SP8 confocal microscope (Leica).

### 4.6 Statistics

Statistical analysis employed Graphpad Prism 9 software. Mann-Whitney U test and Student’s t-test were used to compare means between pairs of groups, and two-way ANOVA was used to compare two groups with two independent variables. A p value less than 0.05 was considered significant. All outliers were included in the analysis.

## Author contributions

**Tatsuya Ishikawa**: Conceptualization, Writing - Original Draft, Investigation, Formal analysis, Funding acquisition, Visualization **Kenta Horie**: Investigation, Formal analysis **Yuki Takakura**: Investigation **Houko Ohki**: Resources **Yuya Maruyama:** Resources, Writing - Review & Editing **Mio Hayama**: Resources, Writing - Review & Editing **Maki Miyauchi**: Resources, Methodology **Takahisa Miyao**: Methodology, Writing - Review & Editing **Naho Hagiwara:** Investigation **Tetsuya J. Kobayashi:** Formal analysis. **Nobuko Akiyama**: Project administration, Writing - Review & Editing, Funding acquisition **Taishin Akiyama**: Conceptualization, Supervision, Writing - Original Draft, Project administration, Funding acquisition

## Acknowledgments

This work was supported by Grants-in-Aid for Scientific Research from JSPS (20K07332, 20H03441) (TA, NA), and CREST from Japan Science and Technology Agency (JPMJCR2011) (TA, TI). TI is supported by the establishment of university fellowships towards the creation of science technology innovation from Japan Science and Technology Agency (JPMJFS2140) and RIKEN Junior Associate Program. Computations were performed on the NIG supercomputer at ROIS, National Institute of Genetics and HOKUSAI supercomputer at ISD, RIKEN.

