## Supplementary Figures for "TCR Repertoire Analysis of CD4-Positive T Cells from Blood and an Affected Organ in an Autoimmune Mouse Model"

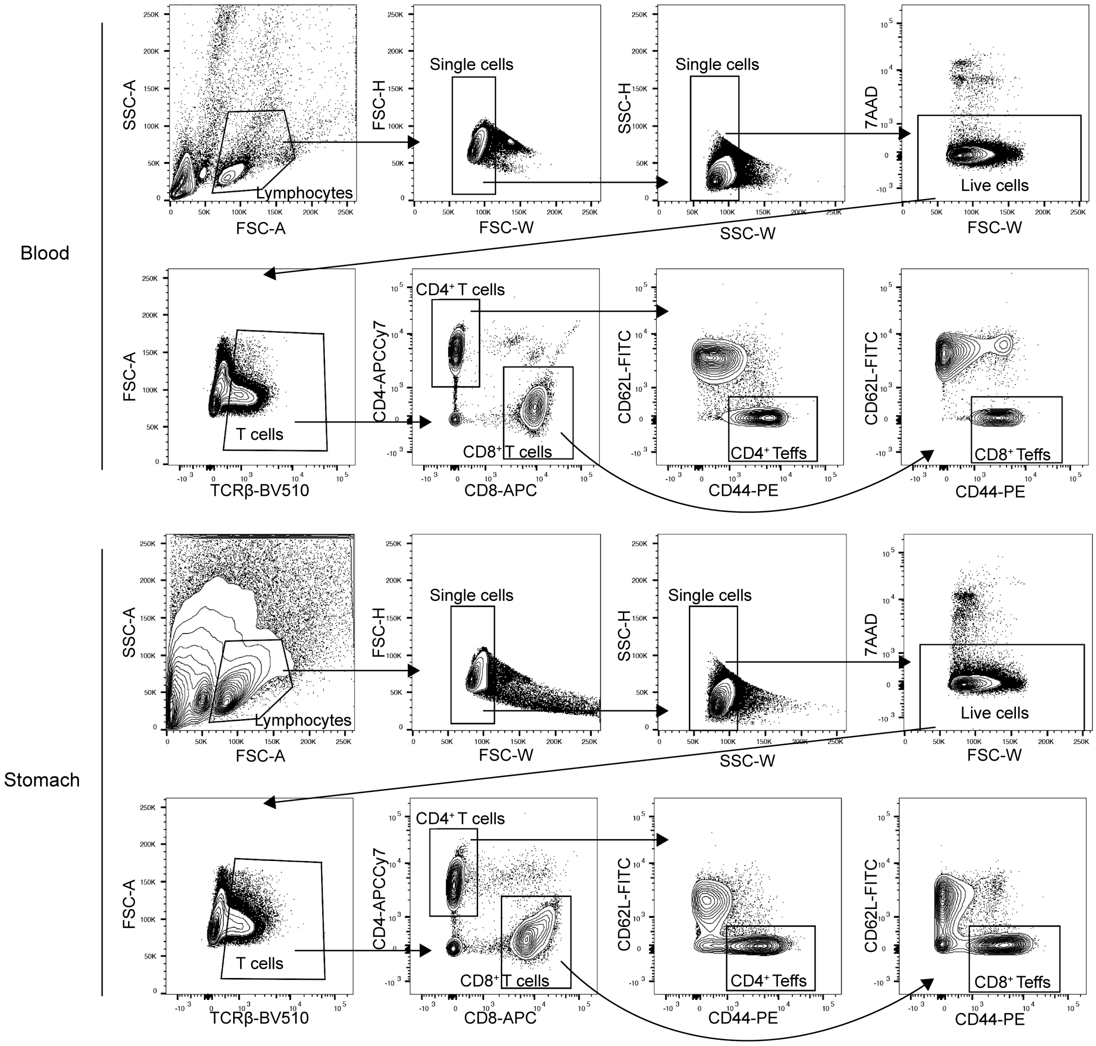


Figure S1

**Gating strategy of Teffs**. a) Representative FACS plots of the gating strategy of CD4/8 T cells in the blood (top) and stomach (bottom).

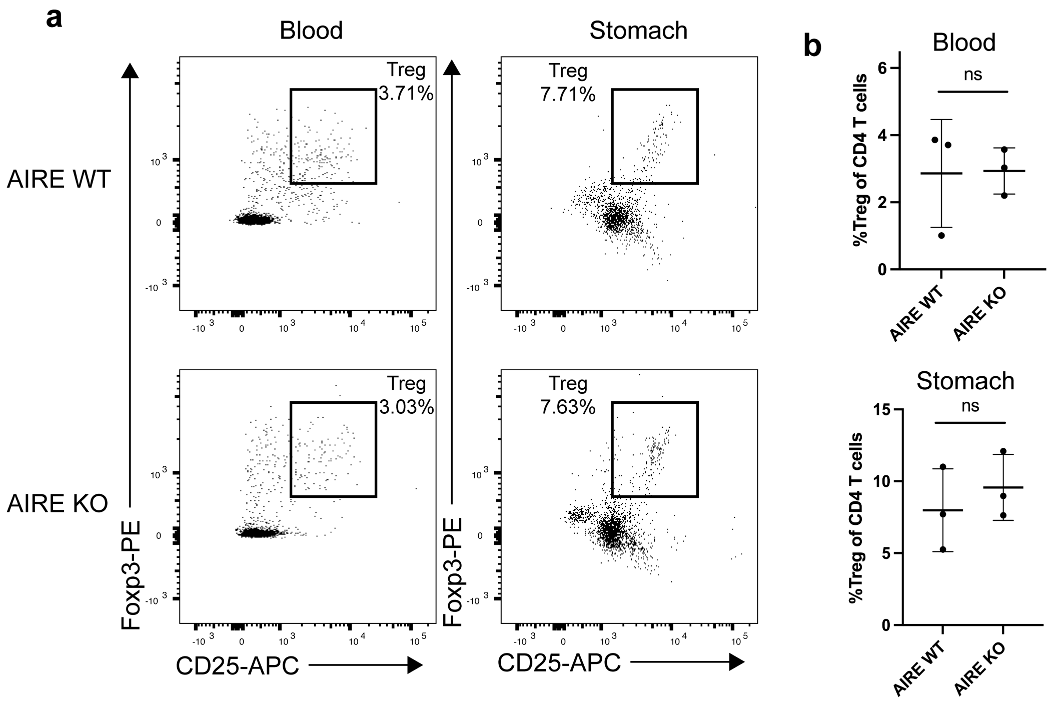


Figure S2

**Detection of Tregs in the blood and stomach**. a) Representative FACS plots showing the detection of Tregs in the blood and stomach of 3-month-old AIRE WT and AIRE KO mice. Plots were gated on $CD4^+$ T cells. b) Quantitative analysis of Tregs from the blood (top) and stomach (bottom). Mann-Whitney U test. ns, not significant ($p > 0.05$).

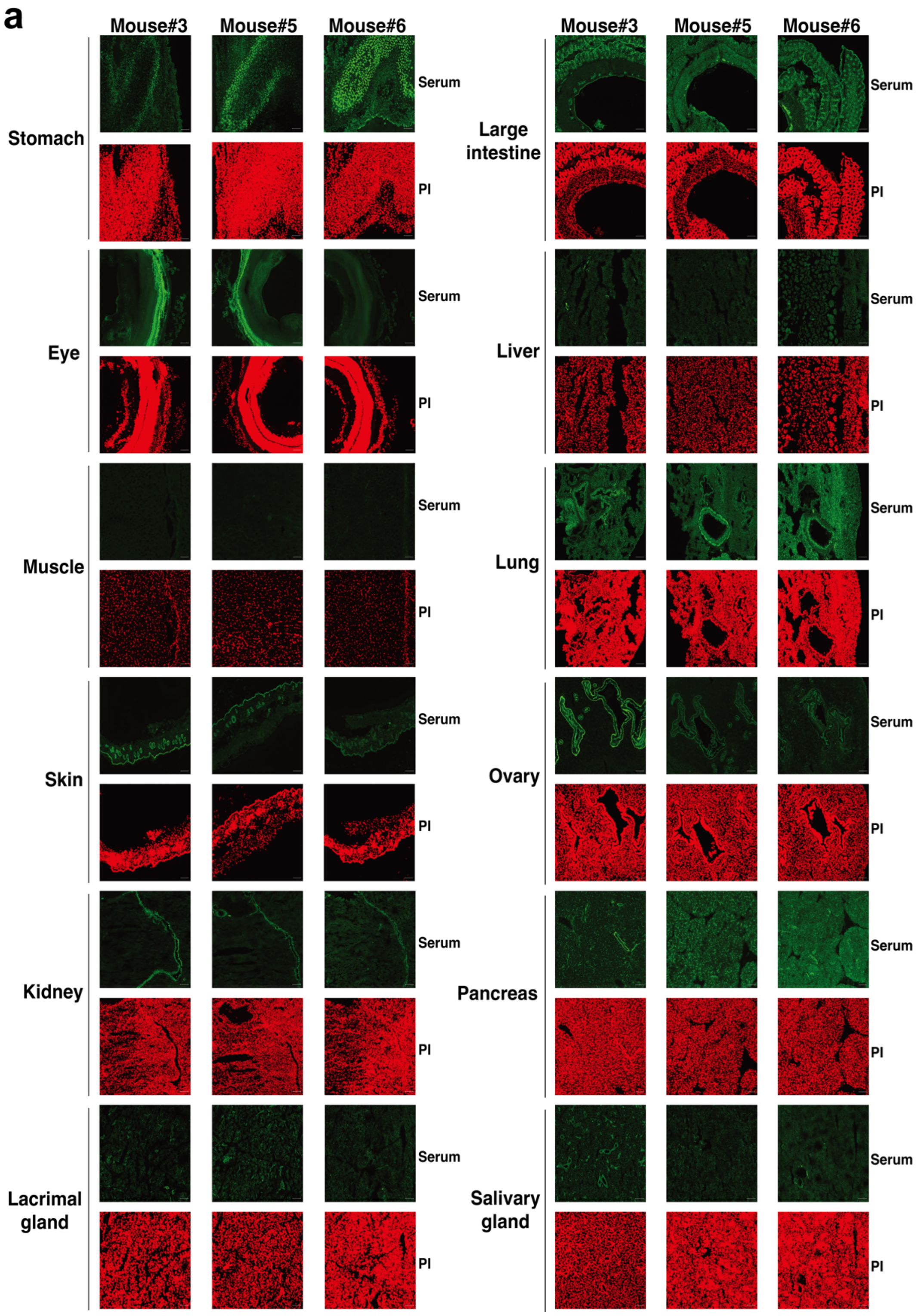


Figure S3

**Immunohistochemistry of sera**. a) Immunofluorescent staining of sera from AIRE KO mice against different tissue sections from Rag1$^{-/-}$ mice.

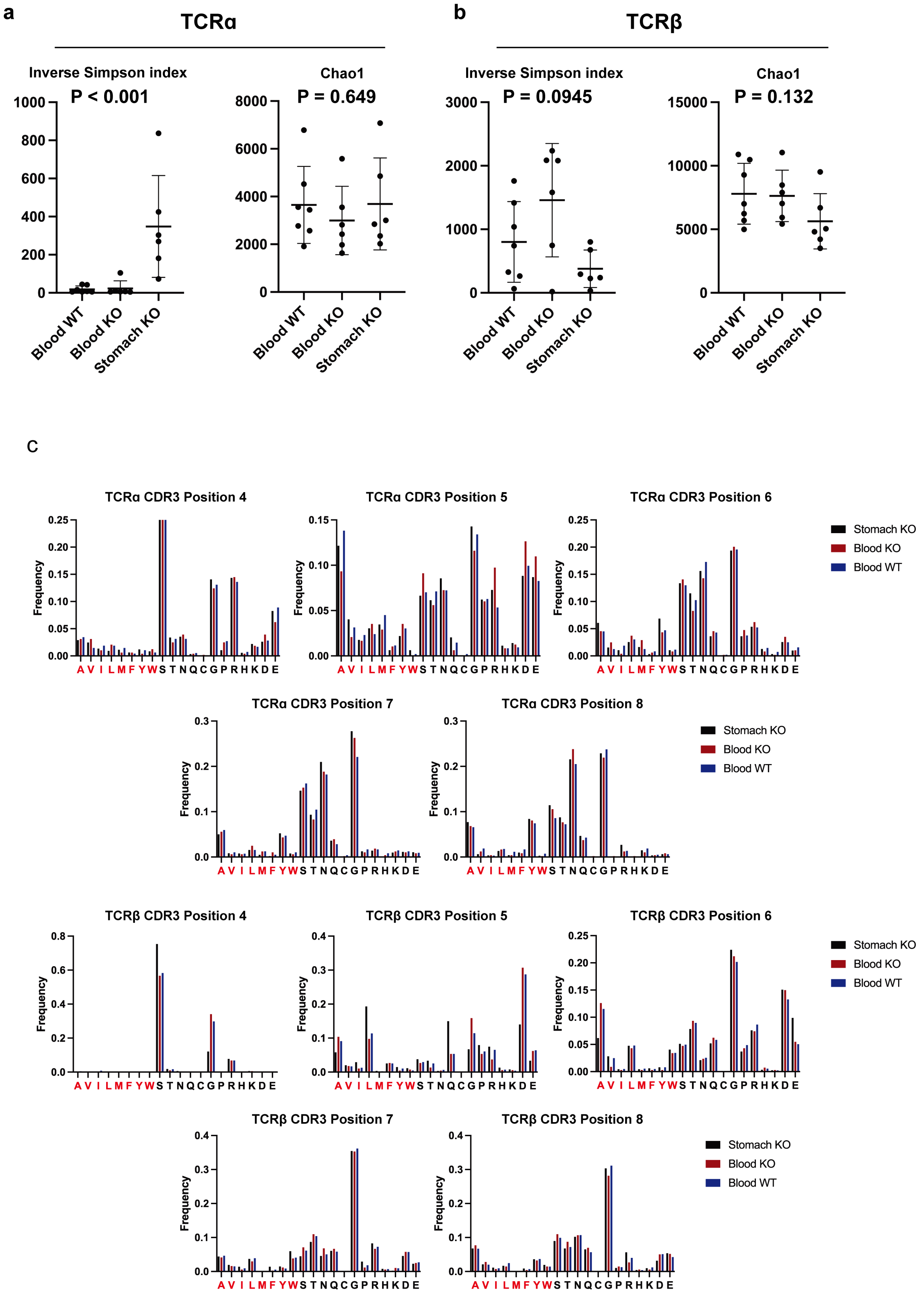


Figure S4

**Diversity metrics for TCRα and β repertoires and amino acid usage of CDR3**.

Inverse Simpson index and Chao1 values of Blood WT, Blood KO, and Stomach KO for a) TCRα and b) TCR β repertoires. Kruskal-Wallis test was used for statistical analyse. c) Frequencies of each amino acid at the position 6 and 7 of CDR3s detected from the stomach of at least two AIRE KO mice (red) and all CDR3s detected from the blood and stomach (blue) are shown for both TCRα and TCRβ.
